# Top-down feedback can explain the existence of working memory traces in early visual cortex

**DOI:** 10.1101/2025.11.27.690959

**Authors:** N. N. Krause, A. Compte, R. L. Rademaker

**Affiliations:** Ernst Strüngmann Institute for Neuroscience in Cooperation with Max Planck Society, Frankfurt am Main, Germany; Vrije Universiteit Amsterdam, Faculty of Behavioural and Movement Sciences, Amsterdam, The Netherlands; Institut d’Investigacions Biomèdiques August Pi i Sunyer (IDIBAPS), Barcelona, Spain

**Author notes:** Correspondence concerning this article should be addressed to Noa N. Krause, Ernst Strüngmann Institute for Neuroscience in Cooperation with Max Planck Society, Frankfurt am Main, Germany.

**Keywords:** working memory, sensory recruitment, neural networks, short-term synaptic plasticity, feedback

## Abstract

The temporary storage of visual memories is known to engage early visual cortex (EVC), where mnemonic content is presumed to be stored in weak or latent states. Two mechanisms have been suggested to underlie this storage: Top-down feedback from anterior cortical sites, and short-term synaptic plasticity (STSP) traces laid down by previous sensory inputs. Here, we tested if these mechanisms can account for two hallmarks of working memory, namely, the flexible selection of information from working memory, and the robustness of working memory against distraction. Simulating firing rates in a neural network model of working memory, we show that STSP cannot easily account for selection of information from within the working memory store, and is not robust to visual distraction. Rather, any input that passes through the visual system is briefly remembered, irrespective of its relevance. In contrast, top-down feedback can establish durable mnemonic signals in EVC that reflect currently task-relevant working memory contents and are robust to distraction. We argue that feedback is a parsimonious explanation for the existence of mnemonic signals in EVC, given that the storage of working memories is distributed across a hierarchy of cortical sites, which are known to have widespread feedback projections to visual cortex.

## Top-down feedback can explain the existence of working memory traces in early visual cortex

Visual working memory (VWM) refers to the ability to temporarily store and manipulate visual information that is perceptually unavailable, and use it to guide behavior [1, 2]. Historically, the short-term retention of visual information has been ascribed to activity in frontal and parietal areas of the brain, such as the dorsolateral prefrontal cortex (dorsolateral PFC), where single neurons fire persistently during the interval between a physical stimulus and a delayed response [3, 4, 5, 6, 7, 8, 9, 10]. Human fMRI studies have similarly shown increases in the blood oxygenation-level dependent (BOLD) response in frontal cortex when visual stimuli are held in mind [11, 12, 13, 14, 15]. Studies in recent decades have moved beyond frontal cortex, and this shift in focus has revealed a widely distributed network of brain areas that support VWM [16, 17, 13, 18, 19, 20] (see [21] for a review).

Among others, the early visual cortex (EVC) has repeatedly been implicated in VWM [22, 23, 24, 25, 26, 27, 28, 29, 30, 12], with fMRI studies consistently reporting decodable VWM signals in EVC response patterns that are sustained for ten or more seconds [24, 25, 31]. While such VWM signals are notably less sustained in human EEG or macaque electrophysiology studies [32, 33, 34, 26, 35] (but see also [23, 29]), they can nevertheless be recovered momentarily by perturbing visual cortex with neutral visual input [26, 35, 36, 37]. The exact neural mechanisms underlying such VWM signals in EVC are under debate, and here we directly compare the plausibility of the two most prominent mechanisms that have been suggested in the literature: Top-down feedback and short-term synaptic plasticity (STSP).

Top-down feedback from frontal and parietal areas to lower-level sensory cortex has long been implicated in visual attention [38, 39, 40, 41, 42], and might also account for weak VWM signals in EVC (e.g., see [43, 44, 45] for reviews). Evidence for this idea comes from the existence of feedback pathways between prefrontal and extrastriate areas [46, 47, 48], and coupling between these areas during VWM [34, 39, 49]. In fact, VWM maintenance predominantly activates the superficial and deep layers of V1 [50, 29], which are strongly innervated by feedback connections [51]. Moreover, VWM signals in EVC are robust to visual distraction or re-emerge after removal of a distractor [20, 31, 27, 29, 52], and VWM signals spread across both hemispheres after being initially encoded in only one hemisphere [53] – such findings are indicative of a sustained VWM source in frontal and parietal areas that can (re-)inject signals into EVC.

More evidence of top-down EVC recruitment comes from studies reporting that activity in visual areas is modulated by VWM contents. For example, work by Huang et al. [54] showed that spontaneous activity in macaque V1 is modulated by the orientation that a monkey holds in mind. This modulation is neither extinguished by distracting visual input, nor is it a mere echo of the coding scheme during stimulus encoding, making top-down feedback a likely source of this modulation. In line with the idea of a sub-threshold modulation in EVC, others have demonstrated that macaque lateral PFC sends feedback signals to the motion-selective middle temporal area, where these signals modulate the local field potential (LFP), but are not strong enough to change spiking activity [34]. These findings point toward latent VWM traces in visual cortex, which manifest as modulations but not as sustained neuronal firing. This aligns with other studies that found the effect of feedback on neural activity in visual cortex to be explicitly gated by bottom-up stimulation, i.e., feedback evoked no activity in visual cortex in the absence of external drive [55, 56, 33] (but see [57, 58, 29]).

STSP is a second contender mechanism by which VWM signals might be maintained in EVC. STSP refers to a well-known mechanism by which the efficacy of synaptic transmission between two neurons is altered by their recent firing history [59, 60]. Specifically, pre-synaptic activity leads to neurotransmitter depletion and calcium accumulation. Neurotransmitter depletion decreases signal throughput, while calcium availability increases signal throughput (e.g., by increasing the probability that vesicles are released from the pre-synaptic terminal [61]). This mechanisms allows activity evoked by an input (e.g., a perceptual stimulus) to temporarily alter a network’s functional connectivity. This altered connectivity pattern can be maintained over a period of seconds and act as a filter, such that newly incoming information propagates more easily through facilitated synapses than through non-facilitated synapses.

Modeling work has shown STSP to be a feasible mechanism of short-term memory retention [60, 62] that can aid the robustness of representations stored in neural networks [63, 64]. While STSP is a brain-area agnostic mechanism, it was initially suggested to support VWM maintenance in PFC: With the idea of an unbroken chain of sustained neural activity coding for VWM content in PFC coming under scrutiny [65, 66], STSP was proposed to carry information across brief periods of silence between bursts of firing [66]. But of course, STSP could just as readily function as a potential memory mechanism in EVC. For example, Pals et al. [67] showed that a neural network model equipped with STSP could encode and retain visual stimuli in latent (i.e., activity-silent) states for several seconds. Perturbation of the neural network led to a reactivation of stimulus-specific activity in a similar manner as observed by Wolff et al. [26] in posterior parts of the brain.

Although there is evidence to suggest that both feedback and STSP might be at play in EVC, the plausibility of each mechanism is still unclear. Can either (or both) explain key signatures of VWM signals in EVC? To investigate this, we tested if the behavior of neural network models equipped with either STSP or feedback agreed with two hallmarks of VWM found in experimental studies: VWM signals in EVC reflect the flexible selection of information, and these signals are robust against visual distraction. Flexible selection of information refers to findings that neural signals in EVC reflect only the currently selected contents held in VWM, rather than a mere perceptual trace. For example, when an item is no longer task-relevant, neural representations for that item are also no longer evident in EVC [26, 68]. Robustness against visual distraction reflects findings that neural traces in EVC are typically not overwritten by competing visual input. Instead, VWM signals either persist or re-emerge after distractor offset [20, 31, 27, 29, 52, 54] (but see [19]).

Here, we presented two tasks to our neural network models to test each of these two hallmarks of VWM storage. First, we tested whether feedback and/or STSP can account for flexible selection by simulating a retro-cue VWM task (as used by Wolff et al. [26]). Specifically, the models were initially tasked with remembering two memory targets until a retro-cue marked only one item as relevant, upon which the other item was expected to be forgotten. Second, we tested whether feedback and/or STSP can account for robustness against distraction by simulating a novel distractor VWM task. The models were tasked with remembering two memory targets, followed by two distractors shown during the delay which were expected to be ignored. In both tasks, we perturbed the models with neural ’impulses’ during the delay to test which information was maintained during different task periods. Such a neural perturbation approach can reveal weak or latent representations (inherent to both the feedback and STSP models) that might not otherwise be detectable in broadband neural activity [35, 26].

Our results show that an STSP mechanism cannot explain flexible selection of information from the VWM store, and is not robust against distraction. Rather, the STSP mechanism lays down a neural trace that reflects any and all information that passes EVC, irrespective of task relevance. In contrast, top-down feedback can account for the mnemonic nature of neural signals in EVC, featuring both flexible selection from the VWM store and robustness against distraction.

## Results

We developed two feedforward neural network models, each comprised of an EVC module and a frontal module, with each module consisting of a left and right hemisphere (Fig 1a-b; see also Materials and Methods). Each hemisphere is a circuit of interconnected excitatory and inhibitory neural populations (Fig. 1a) tuned to produce transient (EVC; Fig. 1c, upper panel) or ring attractor dynamics (frontal; Fig. 1c, lower panel). EVC feeds activity to the frontal module, and left and right frontal circuits mutually inhibit one another. To isolate STSP and feedback mechanisms, one model was equipped with STSP on the EVC module, while the other one was equipped with a feedback connection from the frontal module back to EVC (see Fig. 1b for model architectures and Fig 1d for calcium and neurotransmitter time courses in the STSP model). We simulated firing rates in the EVC and frontal modules of both models in response to the two VWM tasks outlined above (see Fig. 1e for an illustration of how input is presented to the model). To investigate which information was represented in EVC at different times during each task (e.g., in response to memory targets or impulses), we employed a distance-based orientation decoding approach [26].

**Figure 1:**
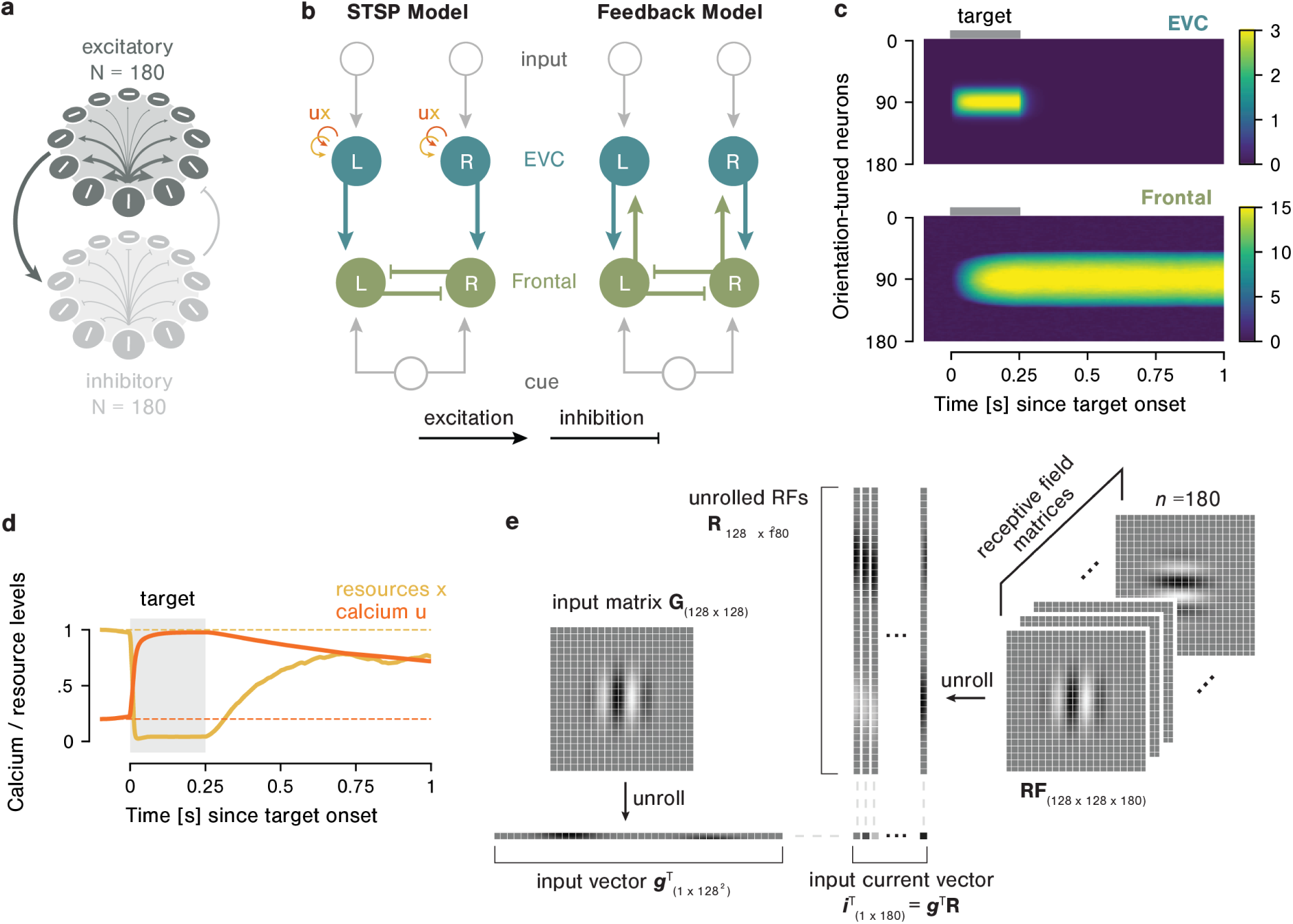
a. Example circuit of interconnected excitatory (E-) and inhibitory (I-) neurons. E-neurons (N=180) have a von Mises-shaped connectivity profile, exciting nearby neighbors more strongly than distant neighbors. Connections among I-neurons (N=180), and between I-and E-neurons, are normalized such that each neuron receives the average activity of the efferent population as its input. **b** Both models consist of an EVC and a frontal module, each comprised of one circuit per hemisphere. Two memory targets (oriented Gabors) are presented separately, one to each EVC hemisphere. A spatial cue can be presented to the frontal module. The models differ in two important ways: The STSP model lacks a feedback connection from frontal to EVC, and is equipped with an STSP mechanism on its EVC circuits. **c** Circuit parameters were tuned to produce transient dynamics in EVC (top) and ring attractor dynamics in frontal modules (bottom). **d** Calcium (u) and neurotransmitter resource (x) dynamics in EVC of the STSP model: In response to neural activity, neurotransmitter resources are depleted while calcium is accumulated. Both variables rebound with individual time courses, which happens faster for neurotransmitter resources than for calcium. **e** Memory targets consist of oriented Gabors (e.g., input matrix **G**) that are presented to each EVC hemisphere, where they are encoded by the neurons’ receptive fields (matrix **RF**). Specifically, the input current vector (**g**^T^ **R**) presented to each EVC hemisphere equals the dot product between unrolled Gabors (**g**^T^) and the unrolled receptive field matrix (**R**).

### Feedback and STSP can explain reactivation of latent traces

As a first step, we verified that both models stored VWM content in weak or latent states in EVC and responded to perturbations with a reactivation of stimulus-specific neural activity (Fig. 2). Specifically, we presented two independently oriented Gabor patches as memory targets, one to each EVC hemisphere, for 250 ms. Models were perturbed during the delay via the presentation of an ’impulse’ (uniform activation of all EVC neurons) for 100 ms.

**Figure 2:**
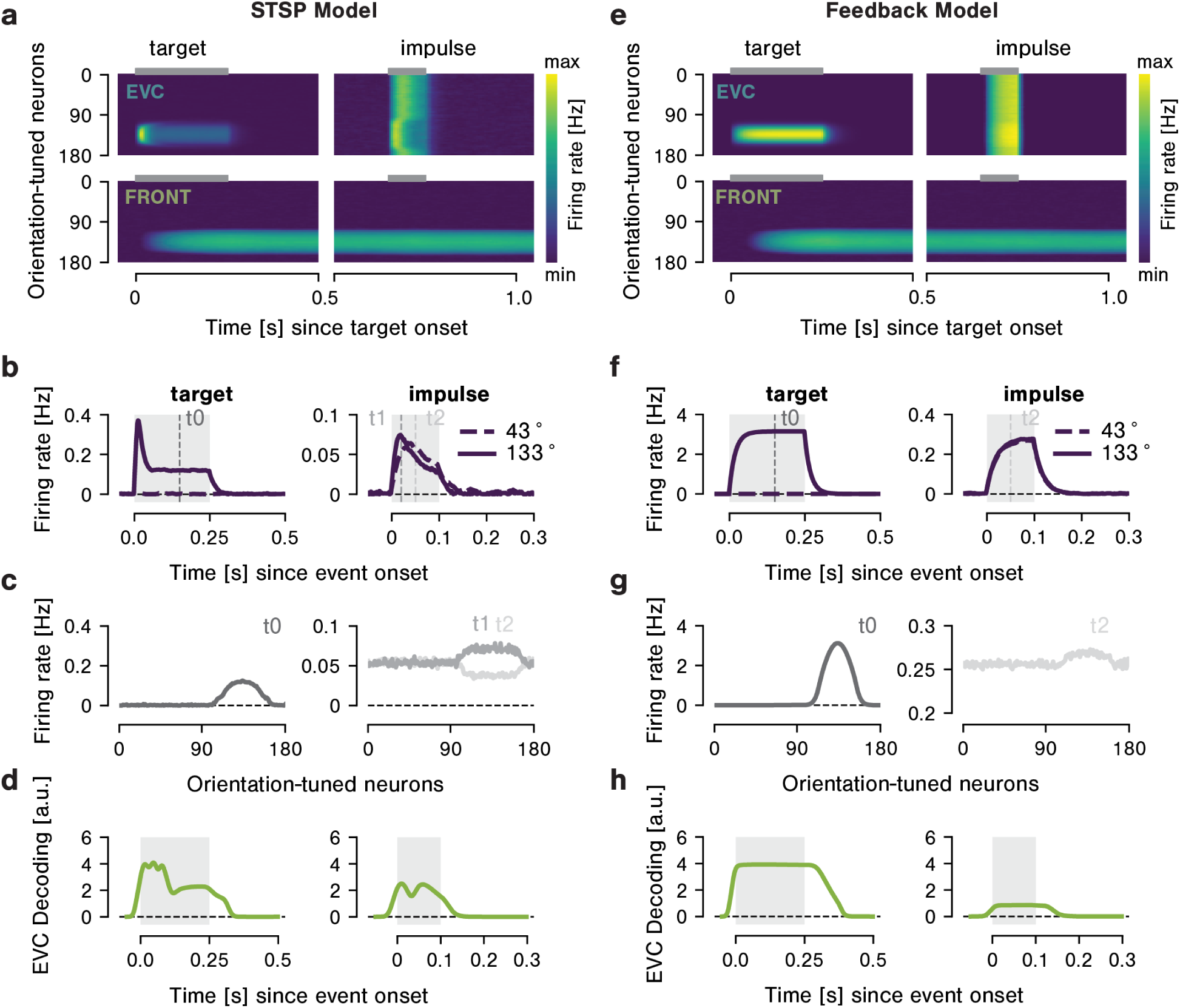
a-d. STSP model. **a** Neural activation in EVC and frontal modules in an example trial and hemisphere. The y-axis shows neurons’ orientation preference; the x-axis shows time since target onset. Color (from purple to yellow) indicates neural firing rate (Hz). On this example trial, a target of 133° was shown. This target evokes a stimulus-specific response that is transient in EVC and sustained in the frontal module. The impulse activates all EVC neurons, but this response differs for neurons that are selective for the target orientation compared to those that have different selectivity. Note that each epoch (target, first impulse) is scaled individually to make the reactivation effect during the impulses visible. **b** Time course of target- (left) and impulse-evoked (right) activity in EVC neurons that are selective (solid line, tuned to 133°) and non-selective (dashed line, tuned to 43°) to the target orientation. The target activates only target-selective neurons (left panel). The impulse activates all neurons, but at time point t1 the target selective neurons respond slightly more than non-selective neurons. This pattern reverses later during the impulse (t2), with lower responses in selective compared to non-selective neurons. **c** Target- and impulse-evoked response profile across all 180 EVC neurons during target (t0, left panel) and impulse presentation (t1 and t2, right panel). The impulse-evoked activity quickly depletes neurotransmitter resources of target-selective neurons, leading to a flip in the population response from t1 to t2. **d** Target orientation can be decoded during target and impulse presentation, but not during the delay period in between. **g-h** Same plots for the Feedback model. Also here, the impulse activates target-selective neurons slightly more than non-selective neurons (**e** and **f**), but there is no pattern reversal during the impulse. Across the entire population, a bump in firing rates is apparent around target-selective neurons (**g**, right panel), which leads to target decoding in response to the impulse (**h**, right panel).

Replicating previous work [67], we show that the STSP model maintains information in a latent state in EVC that can be recovered by perturbations (Fig. 2a-d). In this model, EVC neurons selective for a specific target orientation (e.g., 133°) become activated when that target is shown (Fig. 2a, upper left panel, and time point t0 in Fig. 2b-c, left panels), and the target can be decoded from the population activity (Fig. 2d, left panel). This activation also leaves a trace of facilitation for the target-selective neurons.

Thus, while all EVC neurons fire in response to the impulse (Fig. 2a, upper right panel), previously facilitated neurons respond more vigorously than non-facilitated neurons (time point t1 in Fig. 2b-c, right panels). However, due to the rapid expenditure of neurotransmitter resources in response to the impulse, this pattern reverses after about 30-40 ms, with the previously facilitated neurons now responding less vigorously than the previously non-facilitated neurons (time point t2 in Fig. 2b-c, right panels). Since this response pattern is the inverse of the initial perceptual code (i.e., neurons selective for the target orientation now have systematically lower responses than non-selective neurons), the target orientation can nevertheless be decoded from EVC population activity in the STSP model throughout impulse presentation (Fig. 2d, right panel). Note that while target information is sent to frontal modules, where it is retained in persistent activity (Fig. 2a, bottom panels), this signal cannot propagate back to EVC due to the strict feedforward flow of information in the STSP model. This means that the memory signal in EVC of this model can only be attributed to the STSP mechanism.

The Feedback model can similarly account for weak or latent storage and reactivation effects in EVC (Fig. 2e-h). Following target encoding, neurons in EVC return to baseline firing rates (Fig. 2e, top left panel), but the orientation information is still retained in persistent activity in the frontal modules (Fig. 2e, bottom panels) that project back to EVC. This feedback signal is revealed during the impulse (Fig. 2e, upper right panels), which elicits a slightly stronger response to neurons selective to the target orientation compared to non-selective neurons (Fig. 2f, right panel). Although the reactivation of the target pattern is relatively weak (note the magnified scaling on Fig. 2g, right panel; see suppl. Table A for an overview of scaling in each period), it is consistent across trials. Therefore, target orientation can also be decoded from the EVC of the Feedback model during presentation of both the target and the impulse (Fig. 2h, left and right panels, respectively).

### Feedback, but not STSP, can account for flexible selection from VWM

Presenting the retro-cue task used by Wolff et al. [26] to our models (Fig. 3a), we show that the STSP model cannot flexibly select from among memoranda stored in VWM, while the Feedback model can. Each model initially stores two targets and is expected to select one (and correspondingly, drop the other) in response to a retro-cue, which is modeled as uniform excitatory current added to all neurons of the cued frontal hemisphere (Fig. 1b). Specifically, administration of the retro-cue to one hemisphere shifts the mutual inhibition between left and right frontal circuits in favor of the cued hemisphere, extinguishing activity in the uncued hemisphere (see frontal module activity in Fig. 3b and d).

**Figure 3:**
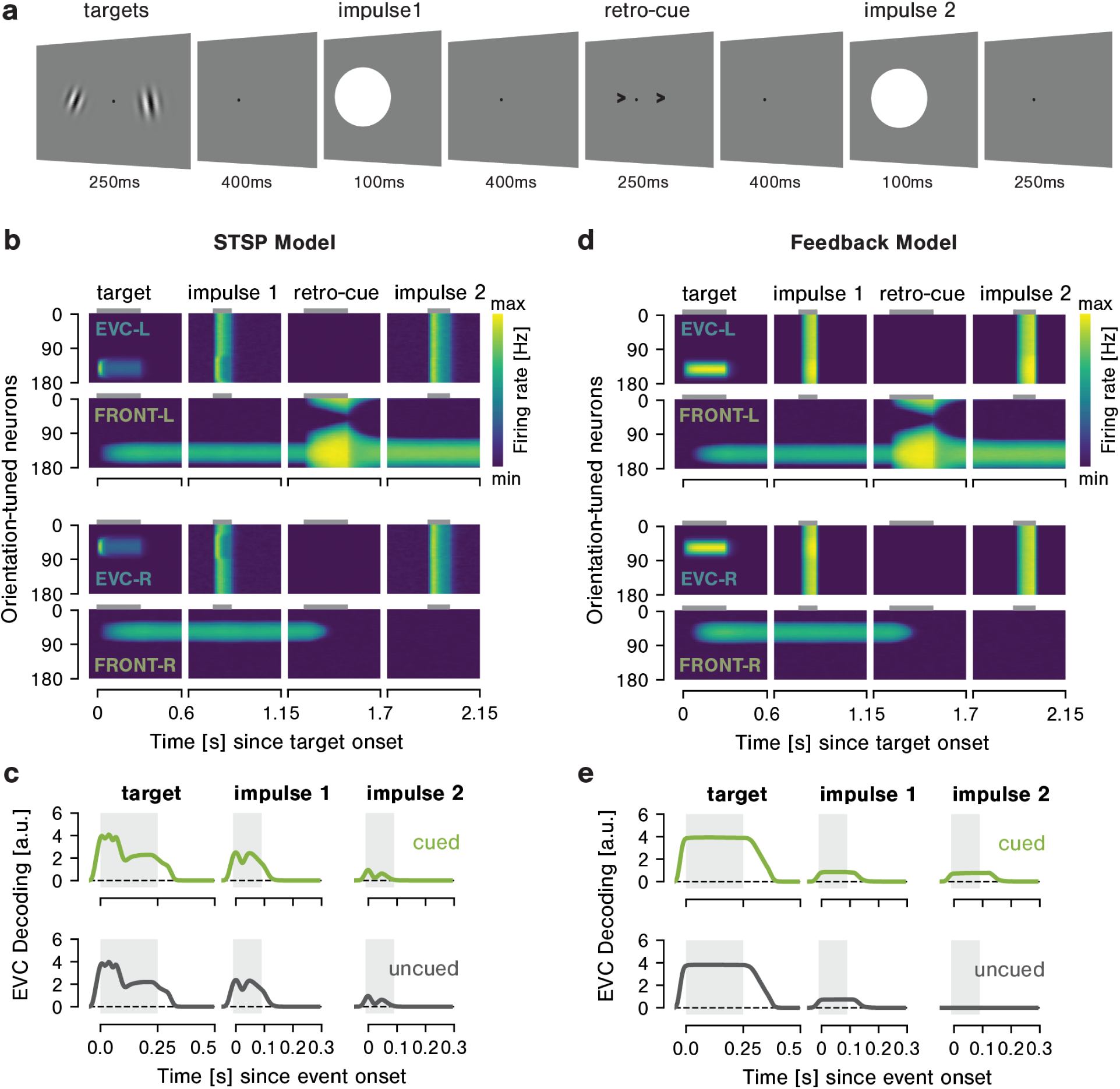
a. Retro-cue task. The model remembers two orientations until a retro-cue (presented to one of the frontal circuits) indicates which target (left or right) is relevant. EVC is perturbed by two impulses during the delay, one before and one after retro-cue presentation. **b** Neural activation in an example trial in the STSP model. In this trial, orientations of 141° and 57° were presented to the left (EVC-L) and right (EVC-R) hemispheres, respectively. EVC panels (1 and 3 from the top): Activity in left (EVC-L) and right (EVC-R) hemispheres throughout all events in the trial. In both EVC hemispheres, transient stimulus-specific activity can be seen in response to the target, and activity across all neurons can be observed in response to each impulse. During the first impulse, responses of target-selective neurons differ from nonselective neurons. While the same is true during the second impulse, the average modification of firing rates is so small (2.4%) that it’s not detectable by eye. Note that each epoch (target, first impulse, distractor, second impulse) is scaled individually to make the reactivation effect during the impulses visible. Frontal panels (2 and 4 from the top): Both left (Front-L) and right (Front-R) frontal hemispheres show sustained stimulus-specific activity before the retro-cue. Administering the retro-cue to one of the frontal hemispheres (here: left) deletes activity in the uncued frontal hemisphere (here: right). **c** Orientation decoding from EVC in response to presentation of memory targets and both impulses. The STSP model does not react to the retro-cue, as both the cued and uncued targets remain decodable at the second impulse. **d-e** Same for the Feedback model with two notable differences in panel **e**: At the second impulse only the cued, but not the uncued, target can be decoded from EVC.

In the STSP model, both cued and uncued target orientations are encoded, and later reactivated by both impulses to EVC (Fig. 3b, EVC panels; note that reactivation during the second impulse, with magnitude of only 2.4 percent, is too weak to be detected visually). Consequently, we were able to decode both orientations from EVC during their presentation (Fig. 3c, left panels) at the first impulse (3c, middle panels), and at the second impulse (Fig. 3c, right panels). Decoding of both cued and uncued targets at the second impulse, ergo after presentation of the retro-cue, indicates that selection did not take place. We purposefully did not equip the STSP model with a feedback connection from frontal to EVC modules in order to isolate feedback vs. STSP as memory mechanisms. However, this also prevented the cue from propagating back to EVC. To allow the cue to reach EVC, we added a feedback connection to the STSP model but gave the frontal module EVC-like tuning in order to avoid ring attractor dynamics (ensuring that the only memory mechanism in the model was still STSP). This architecture did not change our results, as it neither strengthened the cued nor deleted the uncued target representation (suppl. Fig. B1a-b). Arguing that the feedback connection was too weak for the cue to have an effect, we next tried administering the cue directly to EVC, which did the opposite of cueing, and instead weakened or even erased the cued item (suppl. Fig. B1c-d and B2). In sum, none of the cueing strategies we tested here led to selection of the cued item from the VWM store. This indicates that STSP cannot account for flexible selection.

In contrast, the Feedback model *does* reflect the result of flexible selection processes from VWM. To start, EVC encodes and projects information about target orientation to the frontal modules, where persistent stimulus-specific activity emerges (Fig. 3d). This signal is continuously projected back to EVC, but does not cause spiking there outside of impulse epochs. When the retro-cue extinguishes activity in the uncued frontal hemisphere, feedback to EVC from this hemisphere ceases, and only the cued target is decodable from EVC during the second impulse (Fig. 3e). This means that a feedback mechanism can account for flexible selection.

### STSP traces reflect stimulus recency, feedback reflects stimulus relevance

To test whether STSP and feedback mechanisms can explain resistance of VWM signals to distractors, the models were tasked with remembering two targets while ignoring two task-irrelevant distractors (Fig. 4a). We found that STSP cannot withstand the impact of competing irrelevant information during VWM retention, while the Feedback model can.

**Figure 4:**
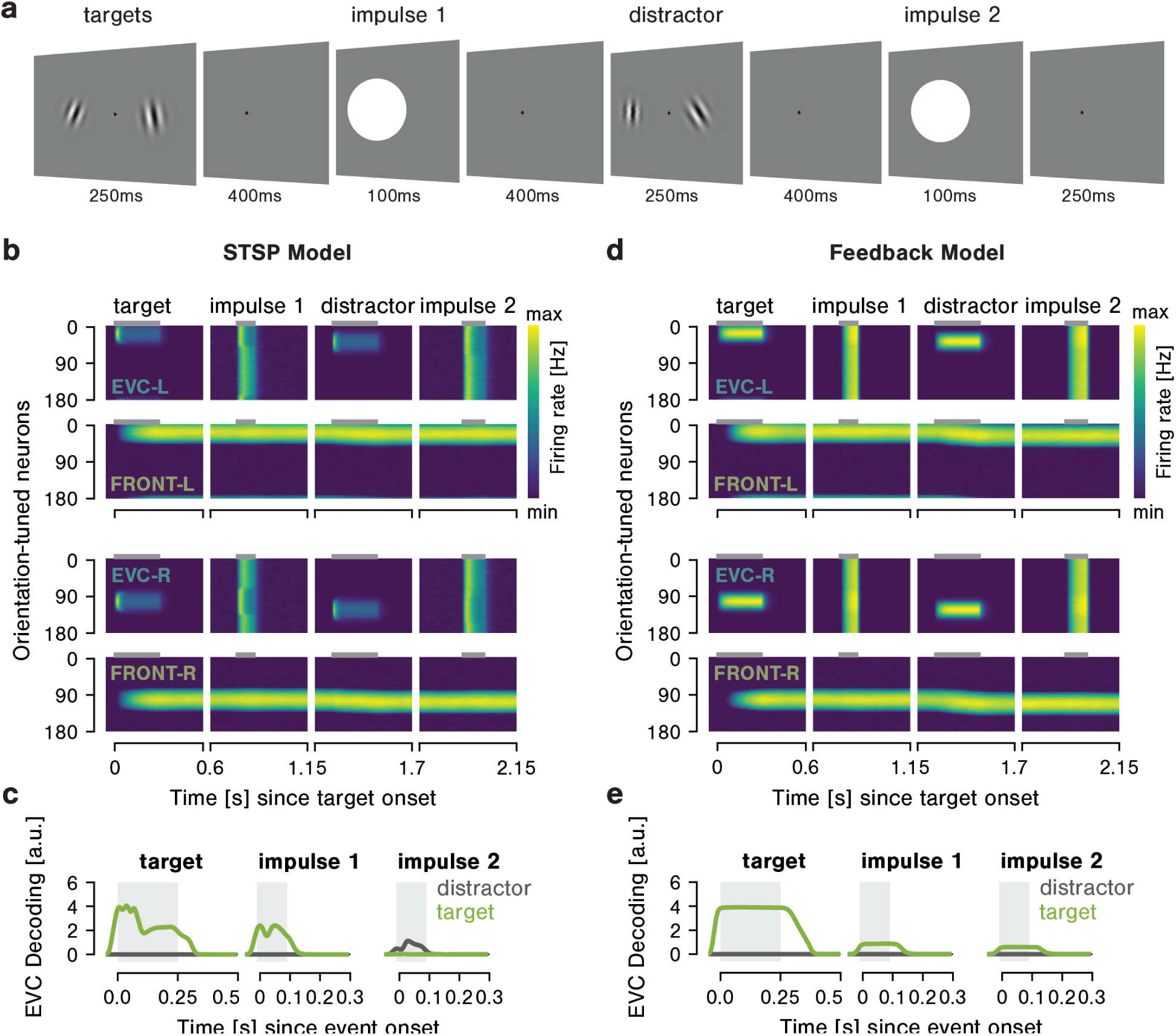
a. Distractor resistance task. The model remembers two orientations while two to-be-ignored distractors are presented to EVC during the delay. Activity is perturbed by two impulses, one shown before, and one shown after distractor presentation. **b** Neural activation time courses in the STSP model for an example trial. In this trial, orientations of 19° and 103° were presented as targets to the left (EVC-L) and right (EVC-R) hemispheres, respectively. Distractors were oriented 20° clockwise of the targets. EVC panels (1 and 3 from the top) show transient stimulus-evoked activity in left and right EVC hemispheres in response to the two target orientations, and activity across all neurons can be observed in response to each impulse. The first impulse activates target-selective neurons differently than non-selective neurons. The second impulse activates only neurons selective to the distractor (though this is again hardly visible, due to the small modulation). Note that each epoch (target, first impulse, distractor, second impulse) is scaled individually to make the reactivation effect during the impulses visible. Frontal panels (2 and 4 from the top) show the left and right frontal hemispheres. Sustained stimulus-specific activity is evident in both hemispheres, and is only slightly affected (biased) by distractor presentation. **c** Orientation decoding from EVC in the STSP model during presentation of the target and both impulses (averaged over left and right hemispheres). The target orientation is decodable during its presentation and during the first impulse. Decoding in response to the second impulse reflects only the distractor, but no longer the target. This indicates that EVC of the STSP model is not robust against distraction. **d-e** Same for the Feedback model with one notable difference in panel **e**: At the second impulse the target is still decodable, whereas the distractor is not. This indicates that EVC of the Feedback model is robust against distraction.

In the STSP model, target orientations were encoded and remembered in EVC during the first half of the task (Fig. 4b), as evidenced by stimulus decoding during their initial presentation (Fig. 4c, left panel) and during the first impulse (Fig. 4c, middle panel). However, presentation of the distractor induced an additional pattern of potentiation in EVC, which was then also maintained. In fact, the second impulse reactivated only the distractor, not the target (Fig. 4c, right panel). Since more time had elapsed since target encoding, calcium and neurotransmitter resources of the initially activated neurons already decayed close to baseline levels, leading to a failure to reactivate the target representation. This means that STSP cannot account for distractor resistance in EVC.

In contrast, the Feedback model does display robustness against visual distraction. Once a target representation has formed in the frontal module, distractors to EVC hardly have an effect on this frontal firing profile and cannot establish a competing representation (Fig. 4d). Since the distractor did not form a new representation in the frontal module, feedback projections to EVC remain target-specific throughout the trial. The target orientation can be decoded during its presentation (Fig. 4e, left panel), the first impulse (Fig. 4e, middle panel), and the second impulse (Fig. 4e, right panel). The distractor could not be decoded from EVC during the second impulse, which was presented after distractor offset (Fig. 4e, right panel). This means that feedback can account for robustness against visual distraction.

### Feedback model predicts attraction toward distractors

While the Feedback model is generally resistant to visual distraction, it should be noted that relatively small target-distractor offsets were associated with attraction of target representations toward distractors (Fig. 5). This attraction bias is evident in EVC (Fig. 5a-b) which inherits it from frontal modules (Fig. 5c-d). This is because frontal modules exhibit attractor dynamics, which cause the target representation to shift slightly in the direction of the distractor once the distractor is presented, and this shift is maintained for the rest of the trial (see Fig. 4d, frontal panels). This shift is propagated back to EVC, where it is revealed during the second impulse (Fig. 5a, red line). We modeled this attraction effect by fitting a first derivative of Gaussian to the bias (circular mean of the population activity vector) for all possible target-distractor offsets, as observed during the second impulse. In frontal modules, results indicate a maximal attraction of 7.38° that peaks for target-distractor offsets around 20° (Fig. 5d), while in EVC the maximal attraction is 6.41° (Fig. 5b). Thus, the Feedforward model predicts that responses to a memory target will be biased toward a task-irrelevant distractor.

**Figure 5:**
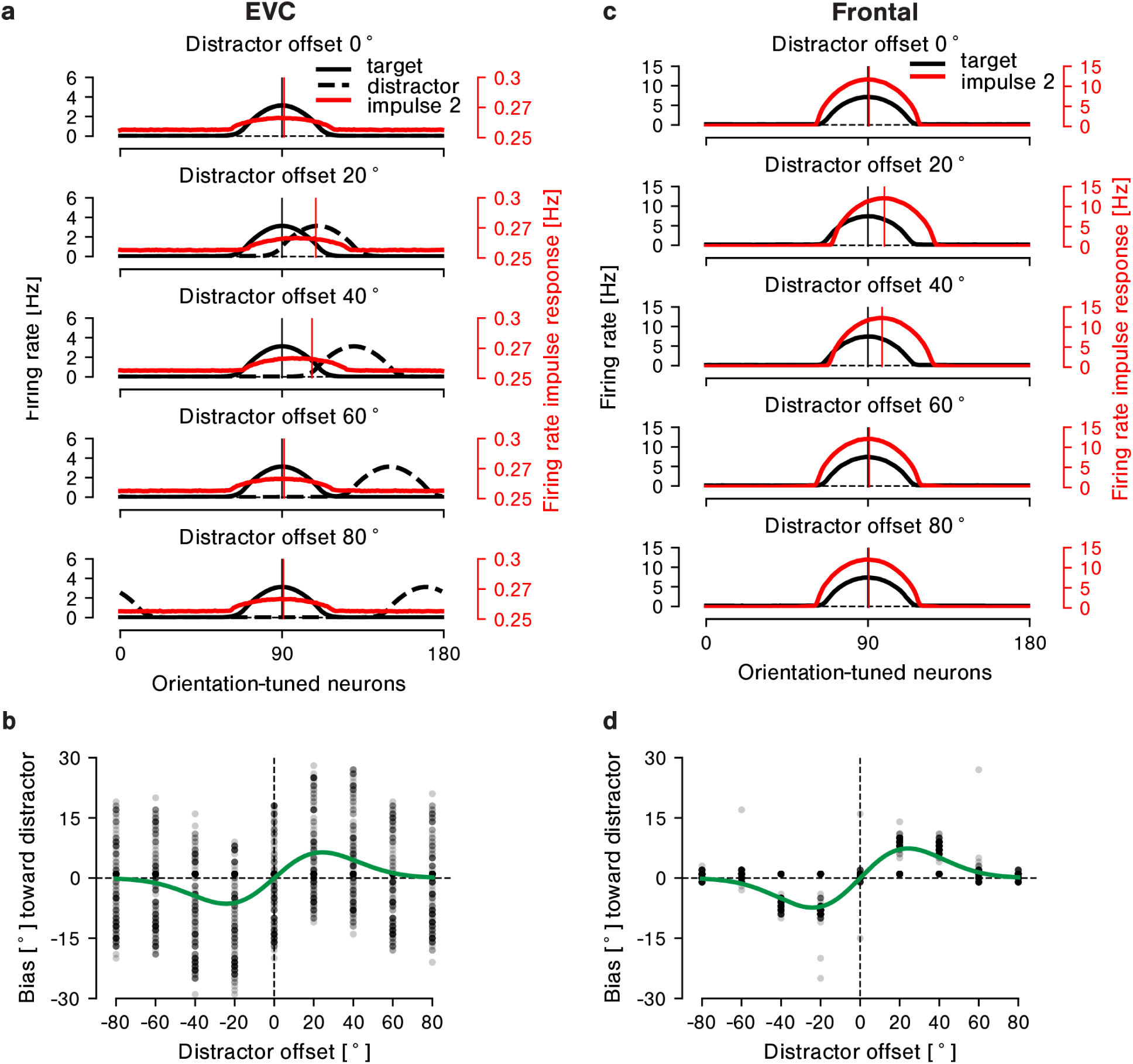
a. EVC responses during presentation of the target (100 ms after onset, black solid line), distractor (100 ms after onset, black dashed line), and second impulse (50 ms after onset, red line with corresponding y-axis shown on the right). From top-to-bottom, responses are shown averaged across all trials where the target-distractor offset was 0°, 20°, 40°, 60°, or 80°. Vertical lines indicate the circular mean of the neural response during target (black) and second impulse (red) presentation, after centering data from all trials on a target orientation of 90°. During the second impulse in the distractor resistance task, target representations in EVC are attracted toward the distractor for target-distractor offsets around 20°-40°. **b** Quantification of target attraction toward the distractor across all trials. The y-axis indicates the shift in degrees relative to the distractor, the x-axis indicates the angular offset between target and distractor. The green line indicates the first derivative of Gaussian fit, while black dots indicate individual trials. The maximal attraction is roughly 7°, which is achieved for target-distractor offsets around 20°. **c-d** The same effects in the frontal module.

## Discussion

The goal of this study was to investigate the merit of two hypotheses regarding how VWM signals are maintained in EVC: top-down feedback and STSP. We know that VWM signals in EVC can manifest as weak modulations of neural firing propensity in visual cortex [34, 54, 66], reflect only task-relevant VWM content [26, 69], and are robust against distraction [20, 31, 70, 52]. To be considered a plausible mechanism for VWM, our two hypotheses should be able to explain such effects. To directly pit feedback and STSP mechanisms against one another, we equipped a neural network model of VWM with either a feedback projection from frontal to visual cortex, or STSP in the visual cortex. We show that both models can maintain VWM signals in a latent "activity-silent" state. We then tested both models on their ability to flexibly select task-relevant items from the VWM store, and their robustness against distraction. We found that while STSP is best thought of as a perceptual buffer that stores an echo of previous visual input, top-down feedback to EVC could explain the presence of mnemonic traces that reflect currently task-relevant VWM content in a distractor-resistant manner.

First, we showed that both feedback and STSP can lead to activity-silent VWM traces in the EVC. For both models, such traces could be uncovered by perturbing EVC with a burst of activity (an ’impulse’). Specifically, this impulse enabled us to decode the VWM target, which was otherwise unobservable during the delay. Importantly, and to our knowledge, this finding establishes for the first time that a feedback mechanism can account for activity-silent storage in EVC equally well as an STSP mechanism – both can account for information maintained in a completely latent state. Having established our basic paradigm and the validity of our two models, we then were able to investigate two hallmarks of VWM representations in EVC derived from experimental studies: Flexible selection of task-relevant information from the VWM store, and the robustness of VWM signals against visual distraction.

To test if the neural traces established in EVC by the two competing models reflect flexible selection from the VWM store, both models were tasked with selecting one of two memory targets on the basis of a retro-cue. Of several strategies attempted to cue the STSP model, none could mimic the pattern of results found by Wolff et al. [26], who could decode only cued (but not uncued) items from posterior brain areas after the retro-cue. Our most straightforward cueing strategy for the STSP model, which was to apply the cue directly to EVC, even had the opposite effect: Increasingly stronger cues weakened and ultimately erased the cued target representation. While we applied several plausible cueing strategies such as this one, we did not exhaust the space of all possible strategies (e.g., inhibitory control via alpha suppression in the hemisphere contralateral to attended objects [71, 72, 73], or top-down attentional gain [74, 75], etc.). We therefore cannot exclude the possibility that more complex strategies might achieve flexible selection in the STSP model – we can only affirm that straightforward cueing strategies did not achieve it. In contrast, the Feedback model replicates the pattern of results found by Wolff et al. [26], and only maintains neural signals for the cued VWM target during the delay. It does so without further adaptations to the cueing strategy. This demonstrates a capacity for flexible selection in the Feedback model, but not in the STSP model.

We also tested whether the neural signals established by feedback and STSP were resistant to visual distraction. The STSP model automatically established a neural trace for any visual input, target or distractor, which is not desirable for a VWM system that requires robustness against distraction. Potentially, an additional ‘shielding’ mechanism, e.g., in the form of top-down attentional gating, could be designed to lessen the impact of distractors on EVC in the STSP model. While such mechanisms are likely at play in cortex [75], this could merely dampen the strength or durability of STSP traces laid down by a distractor. It is unlikely that distractor related signals can be blocked from EVC altogether, considering that distractor suppression only happens at higher levels of the cortical hierarchy, with distractors reaching as far as PFC (albeit with a lesser strength than non-distractors) [40, 76]. The temporally decaying nature of STSP presents an additional challenge, as more recent inputs leave the strongest trace. This is evident from the difference in decoding of targets and distractors during the second impulse – distractors that were presented very recently were decodable, whereas targets presented longer ago were not. Thus, even in case of hypothetical shielding against distraction, STSP is not durable enough for memory storage over intermediate or longer delays. In contrast, neural traces in EVC of the Feedback model reflected robustness against distraction as observed in previous experimental work [20, 31, 70, 52]. Specifically, memory targets remained decodable even after distractor presentation. This demonstrates distractor resistance in the Feedback model, but not in the STSP model.

In the Feedback model, distractor resistance is inherited from higher-level cortex, which we chose to model using a ring attractor [77, 78]. Ring (or bump) attractors have a long-standing history in modeling prefrontal involvement in VWM [77, 78, 79, 80, 81, 82, 83, 84]. Compte et al. [77] already demonstrated that representations in ring attractors can withstand distracting input, and that distractors can shift target representations in their direction, with the magnitude of such a bias depending on target-distractor offset. We replicate these effects in the frontal modules of our model, and extend this finding by showing that attractive biases propagate back to EVC. Attraction of a memory target toward a task-irrelevant distractor is a well-known phenomenon, shown in previous behavioral studies [85, 86, 87, 88, 89], and even in EEG studies looking at posterior parts of the brain [79]. Interestingly, a recent EEG study found such attraction biases when target and distractor were presented at the same location, but found a repulsion when target and distractor were presented on opposite sides of the screen [79]. The authors modeled these effects with a neural network model of similar architecture as our own. Crucially, to explain the repulsion effects found in their EEG data, they added cross-hemispheric inhibition between visual cortex modules. This shows that, with relatively simple changes to its architecture, our model could be adapted to capture increasingly more complex neural behavior in future work.

Note that the current implementation of our Feedback model requires task-relevant information to be reliably retained in a spiking code outside of EVC – we assume in frontal cortex. However, the idea of persistent population activity in higher-order cortex has come under scrutiny in light of sparse and bursty neuronal firing patterns (see [65] for a review) and dynamic changes in the population response [90, 66, 91, 84]. It has been suggested that PFC might maintain information latent states [66]. Future models could explore combining feedback and STSP into the frontal modules of a single model, where STSP can act as a short-term buffer to overcome brief periods of rest in between bursts of prefrontal neuronal activity. Similarly, dynamic coding could be introduced to the model by e.g., random connections between visual and frontal cortex, or by using asymmetric connectivity within the ring attractor [84].

In addition to the nature of memory storage (persistent spiking, dynamic population responses, or latent states), another topic of ongoing debate is whether frontal cortex, and dorsolateral PFC specifically, functions truly as a memory site, or just provides top-down control over other memory sites [21, 92, 93, 94, 95]. While many studies report durable VWM representations in PFC [96, 84, 97, 34, 98, 16] (but see [99]), other sources highlight PFC’s role in exerting attentional control over other areas [100, 93]. For example, VWM signals in the frontal eye fields (e.g., [101]), posterior parietal cortex (e.g., [19, 16, 17]), and several other sites (see [21] for an overview) might rely on top-down control from PFC. The feedback mechanism in our model is largely agnostic to the exact exact cortical area in which VWM contents are stored. The main prerequisite is the existence of feedback connections to EVC, which are known to be widespread in cortex [46, 47, 48, 39, 56, 102, 103]. VWM signals in EVC of our two models were truly activity-silent during the delay, meaning that the remembered information was not reflected in spiking activity. The question whether this is also the case in cortex is still up for debate. On the one hand, several studies report no persistent elevated delay-period activity in macaque visual cortex [32, 34], and cannot decode VWM content from posterior EEG channels outside of a perturbation by an impulse [35, 26], which implies an activity-silent state. On the other hand, several other studies have shown that EVC can exhibit elevated neural activity [54, 29, 23] as well as delay-period decoding from EEG [104, 68, 105, 106], which implies (weak) activity during the delay. In neurophysiology studies, modulated firing rates in EVC often take place in a low firing regime (i.e., a modulation of background firing rates, e.g., [29, 54]), which means these effects require relatively more experimental power to uncover. Especially using EEG, a modulation of baseline firing rates might be difficult to detect given that baseline firing within a population tends to be temporally uncoordinated. Evoked responses to external stimuli can temporally align large populations of neurons and make them more detectable with EEG. This could explain why delay-period decoding can be absent from posterior EEG electrodes outside of periods with evoked responses (e.g., by an impulse).

Relating this back to our models, the efficacy of neither the Feedback nor the STSP model hinges on whether VWM traces in EVC are completely ’activity-silent’. In the STSP model, it’s been shown that an increase in background activity can lead to a bistable regime with spontaneous reactivations of the stored pattern, such that the model oscillates between ’on’ and ’off’ states (states in which the population activity does or does not reflect the stored pattern, respectively) [60, 67]. In the Feedback model, top-down signals arriving in EVC are able to pattern spontaneous activity, as well as activity evoked by external stimuli, and therefore could explain the resurgence of VWM content in response to neural perturbations as well as the modulation of baseline firing rates sometimes reported in macaque visual areas during VWM.

Importantly, a feedback mechanism could also explain another notable discrepancy in the literature, namely, the discrepancy between fleeting decoding of VWM contents from EEG signals [26, 107, 36] versus far more durable decoding from the BOLD signal measured with fMRI [25, 20, 24, 30, 12]. As alluded to above, EEG mainly picks up on highly synchronized activity of large populations of neurons. Diffuse and potentially prolonged top-down modulation might not arrive with the required strength and temporal synchronization to become detectable in the EEG signal. In contrast, the BOLD signal reflects the total input to a population irrespective of whether this input exceeds neural firing thresholds, and might therefore be more sensitive to information arriving via feedback [108].

As is true for all models, we acknowledge that our proposed models are a simplification of more complex processes in cortex. Yet, they provide a mechanistic tool to demonstrate how activity-silent storage in EVC may come about, and can be used to test important hallmarks of VWM (i.e., flexible selection and distractor resistance). In addition to satisfying these hallmarks, our Feedback model can in theory account for other phenomena in the literature, such as patterned EVC responses both with or without truly activity-silent, or the discrepancy between EEG and fMRI findings.

## Conclusion

In summary, we provide proof-of-principle evidence that mnemonic neural traces in EVC can be explained by top-down feedback during VWM maintenance. These feedback signals make mnemonic content available in EVC, possibly acting as a perceptual filter and allowing for interactions between memory content and incoming visual information. While passive mechanisms like STSP also theoretically allow for the integration of stored content and incoming visual information, we demonstrate that STSP alone cannot explain the mnemonic nature of neural traces recovered from EVC.

## Materials and Methods

### Architecture

Models consisted of an EVC and a frontal module, with each module comprised of one circuit per hemisphere (Fig. 1a-b). All circuits had identical architecture but were equipped with varying connection weights to change the dynamics of the circuit. As a rule, the EVC module was tuned to produce transient responses (Fig. 1c, top panel) to input in order to mimic activity profiles observed in EVC in non-human primates [32, 34]. Frontal circuits were tuned to produce ring attractor dynamics [77] (Fig. 1c, bottom panel). Parameter values for the frontal module were based on Wimmer et al. [78], and the same values were scaled by a factor of 0.01 to model EVC (see supplementary Table C1 for an overview over parameter tuning).

Each circuit was comprised of *N* =180 excitatory (E) neurons and *N* =180 inhibitory (I) neurons. E-neurons recurrently excited themselves via all-to-all connectivity matrix **W***_EE_* and excited I-neurons via **W***_EI_*. I-neurons in turn inhibited E-neurons via **W***_IE_* and themselves via **W***_II_*. **W***_EE_* was a circulant matrix comprised of a von Mises (*µ* = 0, *κ* = 5) that mimicked the characteristic local excitation pattern of ring attractor models [78, 77], where E-neurons excite their direct neighbors in feature space strongly while exciting their more distant neighbors weakly or not at all. Matrices **W***_EI_*, **W***_IE_*, and **W***_II_* are all-to-all connectivity matrices normalized by *N*, such that each neuron in the afferent population receives the average activity of the efferent population. Each connectivity matrix is scaled by its own respective constant *g* that controls population activity dynamics. We calculated the input to E-neurons **i***^E^* and I-neurons **i***^I^* as a summation of excitatory and inhibitory input:

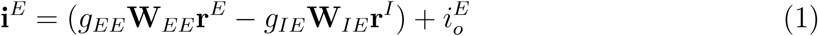

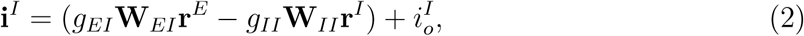

where *g_EE_*, *g_EI_*, *g_IE_*, and *g_II_* scale the connection strength between neural populations within a circuit, **r***^E^* and **r***^I^* represent the vector of firing rates in the E- and I-neurons, respectively, and 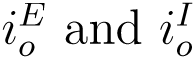 refer to the summed input current arriving at the E-and I-neurons from other cortical sources that were not included in our models. Firing rates were calculated by passing input current through a non-linear function *ϕ*(**i**) [109] and adding Gaussian noise *ξ*(*t*) with SD = *σ*:

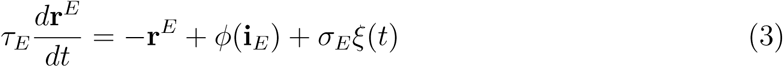

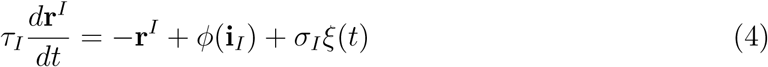

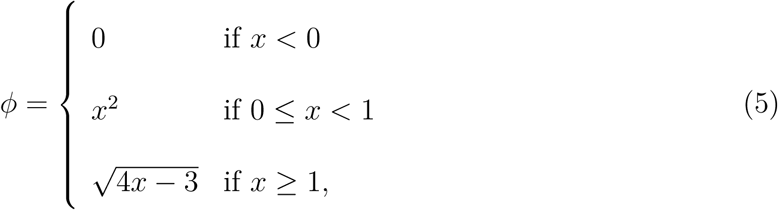

where *τ_E_* and *τ_I_* refer to the synaptic time constants of excitatory and inhibitory neurons, respectively.

E-neurons were tuned to orientation, responding preferentially to some orientations over others. In EVC, neurons’ orientation selectivity arose through their receptive fields (each RF 128 x 128 pixels), which were modeled as Gabor patches (sine-wave gratings of unique integer orientation between 0-180°, phase *π*, convolved with a 2-dimensional Gaussian kernel, *SD* = 1) and act as a spatial filter of the input. Inputs to EVC also consisted of Gabor patches identical to RFs in size, phase, and spatial frequency, but varying in orientation. Input to each E-neuron in response to an oriented grating was calculated as the dot product between the unrolled input and each neuron’s unrolled RF matrix (Fig. 1e). As a result, E-neurons responded maximally to inputs aligned with their RF’s orientation, and responses gradually dropped as the angular distance between input and RF orientations increased. Each EVC hemisphere received separate input and the circuits were not connected to one another.

EVC E-neurons were connected to frontal E-neurons via one-to-one feedforward connections. In addition, left and right frontal circuits mutually inhibited one another in an all-to-all manner: left I-neurons inhibited right E-neurons and vice versa. To model the input to frontal E-neurons, we modified formula (1) by adding an extra term for the feedforward input coming from EVC E-neurons, and for the mutual inhibition coming from frontal I-neurons of the other hemisphere. For example, input current to E-neurons in the left frontal hemisphere is calculated as follows:

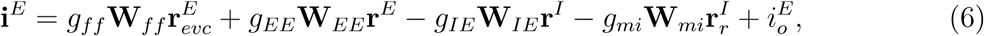

where *g_ff_* determines the feedforward connection strength between EVC E-neurons and frontal E-neurons, **W***_ff_* determines the one-to-one connectivity structure (identity matrix normalized by *N*) between the same populations, and **r***^E^*_evc_ refers to the vector of firing rates of EVC E-neurons. Similarly, *g_mi_* scales the mutual inhibition strength between the left and right frontal circuits, **W***_mi_* determines the all-to-all connectivity structure (normalized by *N*, such that each neuron receives the average activity of the efferent population), and **r***^I^* refers to the vector of firing rates of I-neurons in the right frontal circuit. The same logic was applied to calculate activity in the right frontal hemisphere. Firing rates in the frontal circuits were calculated by passing **i***_E_* and **i***_I_* to formulas (3) and (4), respectively.

### STSP Model

In the STSP model, momentary firing rates of E-neurons were additionally affected by their recent firing history. Specifically, neural activity depressed subsequent synaptic efficacy by depleting neurotransmitter resources (reducing firing rates), but simultaneously facilitated subsequent synaptic efficacy through the accumulation of presynaptic calcium (increasing firing rates). At rest, a neuron had full neurotransmitter stores (*x*_0_ = 1), as well as a baseline probability of releasing neurotransmitters (*u*_0_ = 0.2). Both neurotransmitter levels and calcium decayed back to baseline over time with time constants *τ_D_* = 0.2 and *τ_F_* = 1.5, modeling the decay rate of the depressive effect of neurotransmitter depletion and facilitatory effect of calcium accumulation, respectively (Fig. 1d).

Using kinetic differential equations governing neurotransmitter depletion and calcium accumulation originally formulated by [59], we calculated the momentary levels of neurotransmitter resources *x* and calcium *u* on the basis of the momentary firing rates.

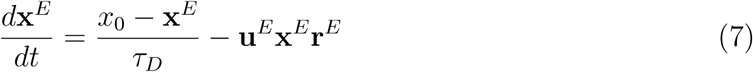

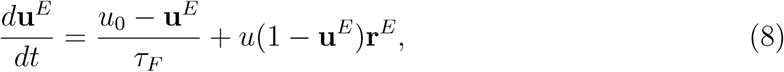

The impact of neurotransmitter and calcium levels on synaptic efficacy was modeled by scaling the firing rate of each EVC E-neuron by the factor **ux** in the calculation of the input current (equation (1)).

### Feedback Model

As an alternative memory mechanism to STSP, we implemented a feedback mechanism. Specifically, E-neurons in the frontal circuits, which retained the memory over time in an attractor state, were connected to E-neurons in the EVC circuits via excitatory feedback connections. Feedback connection strength was set to a low number to achieve latent modulation of neural activity in EVC E-neurons. Thus, we modified formula (1) for the EVC E-neurons in the Feedback model as follows:

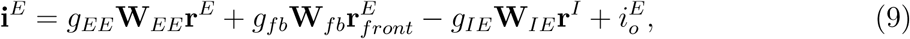

where *g_fb_* and **W***_fb_* represent the strength and structure, respectively, of the feedback connection from frontal E-neurons to EVC E-neurons. The connectivity structure mirrored **W***_ff_* (one-to-one, i.e., identity matrix normalized by *N*). An overview over connections within each model can be found in supplementary Table D1. All models were simulated using custom code run in Python 3.10.

## Simulations

### Flexible Selection Task

To test whether the models could flexibly select and drop information from the VWM store, we presented them with the retro-cue VWM task originally used by Wolff et al. [26] (see Fig. 3a). The models were presented with two Gabor patches (one to each hemisphere) of differing orientations for 250 ms, followed by an initial delay period of 900 ms during which both orientations needed to be remembered. Midway though this first delay, a visual ’impulse’ was presented for 100 ms, which we modeled as a uniform input added to E-neurons of the EVC module (see strength of all inputs in supplementary Table D1). Note that in the original task design with human participants, this impulse was a white circle or a bull’s eye stimulus [26, 35]. Subsequently, a 250 ms retro-cue marked the left or right orientation as the cued target. This retro-cue was modeled as a strong uniform input added to E-neurons in the frontal circuit corresponding to the cued hemisphere. Next, the second part of the delay period of 400 ms ensued, during which only the cued orientation still needed to be remembered, followed by another 100 ms impulse. We sampled cued and uncued orientations randomly from within nine orientation bins (centered on 0°, 20°, 40°, 60°, 80°, 100°, 120°, 140°, 160°), and counter-balanced target with non-target bins and target location (left vs. right). Repeating each unique combination nine times resulted in 1458 trials.

### Distractor Resistance Task

To test whether the models could protect VWM content from visual distraction, we presented models with a distractor VWM task (in the gist of [85]; see Fig. 4a). Models were presented with two Gabor patches (one to each hemisphere) of different orientations for 250 ms, followed by an initial delay period of 900 ms during which both orientations needed to be remembered. Midway though this first delay, a visual impulse was presented for 100 ms. Subsequently, each hemisphere was presented with a to-be-ignored distractor orientation for 250 ms. Next, a second delay period of 400 ms ensued, during which both initial target orientations still needed to be remembered, followed by another 100 ms impulse. The target orientations were sampled from the same orientation bins as for the flexible selection task, and left and right target bins were counterbalanced with 9 possible distractor offsets (0, *±*20, *±*40, *±*60, *±*80). Repeating each unique combination twice resulted in 1458 trials.

## Data Analysis

### Time-Resolved Decoder

To maximize comparability of our modeling results with EEG results by Wolff et al. [26], we used the authors’ freely available code for orientation decoding. The approach assumes that different orientations elicit unique but overlapping neural activation patterns, such that trials with *similar* target orientations are close to each other in high-dimensional neural state space, while trials with dissimilar target orientations are distant from each other. This distance between neural activation patterns (in the firing rates of E-neurons) between trials of varying orientations was quantified using the Mahalanobis distance [110] in a 9-fold cross-validation loop. Briefly, in every fold we randomly picked <u>^1^</u> trials as test data, while the remaining 8/9 of trials were used as training data. Training trials were binned into 9 orientation bins, sub-sampled to ensure equal number of trials per bin, after which we averaged across the trials within each bin. The data was then smoothed across bins by replacing the neural activation pattern in each bin by a weighted sum of the patterns across all bins – the weights for each bin were given by a cosine centered on that bin. Next, the Mahalanobis distance between each bin and each test trial was calculated, resulting in 9 Mahalanobis distances per test trial. The cosine similarity of this resulting distance vector acted as an index of decoding accuracy. Each time-point within a trial was decoded separately, and we repeated the entire cross-validation procedure ten times with new random splits into train and test trials. A full explanation of the approach can be found in [26]. Of note, before submitting neural firing rates to this decoding algorithm, we smoothed the data with a gaussian kernel among the time dimension (*σ* = 8ms) and mean-centered by subtracting the average neural firing rate separately per trial and time point.

### Quantifying target-distractor attraction

To test if distractors had an attractive effect on VWM targets, we tracked how target representations shifted after distractor presentation as a function of target-distractor offset in EVC and in frontal modules. A target representation resembles a bump in the population response that should be roughly centered at the target orientation. To see if this bump was shifted toward the distractor orientation at the time of the second impulse (50 ms after impulse onset), we calculated the circular mean of the population response on every trial, and plotted it against the angular offset between target and distractor. The circular mean of the population response was found by scaling each neuron’s preferred angle by that neuron’s current firing rate *r* (as in [78]):

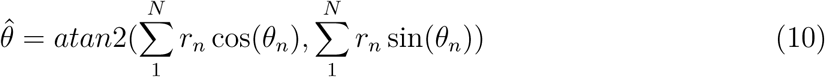

To quantify if there was attraction or repulsion between targets and distractors, we fit the resulting vector of circular means with a first derivative of Gaussian defined as *y* = *xawce^−^*^(^*^wx^*^)2^, where 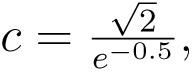 *x* is the orientation of the distractor relative to the target, *a* refers to the curve amplitude, and *w* refers to the curve width [111]. Gaussian derivatives were fit using the scipy.optimize package [112].

## Acknowledgements

We would like to thank NNK’s thesis advisory committee for many helpful comments on this project over the years, and Michael J. Wolff for insightful discussions. RLR and NNK are funded by the Max Planck Society. No AI tools were used in writing the code or manuscript for this project.

## Code Availability

The code used for simulating the models and creating figures is available under https://github.com/NoaNoelle/Feedback_can_explain_EVC_memory_traces.

## Appendix A

### Firing Rate Scaling

Though neurons in EVC respond to perturbation by reactivating the stored neural representation, this effect is small. To make the reactivation effect in the STSP and Feedback models visually detectable, we scaled firing rate panels in Fig. 2a,e, Fig. 3b,d, and Fig. 4b,d differently for different task periods (stimulus vs. impulse presentation). Minimum and maxiumum firing rate values used in each firing rate panel are detailed below.

**Table A1:** Firing rate scaling for different models, modules, and task periods.

| Task Period | STSP Model |  | Feedback Model |  |
| --- | --- | --- | --- | --- |
|  | EVC | Frontal | EVC | Frontal |
| target/retro-cue/distractor | [0, 0.38] | [0, 19] | [0, 3.2] | [0, 19] |
| impulse 1/2 | [0, 0.08] | [0, 19] | [0.1, 0.28] | [0, 19] |

## Appendix B

### Alternative Cueing Strategies for STSP Model

To test whether a similar cueing mechanism as applied to the feedback model might work for the STSP model, we tested different model architectures. First, we added a feedback connection from the frontal modules to the EVC modules, as done in the feedback models. To still isolate the different memory mechanisms (STSP vs. top-down feedback), we introduced EVC-like tuning in the frontal module to avoid memory maintenance in that module. With the intact feedback connection, the cue that was applied to the frontal module could propagate back to EVC. As can be seen in Figure B1a, the cue had no noticeable bearing on activity in EVC. As a result, the second impulse still revealed a neural trace for the both the cued and the uncued items (Figure B1b). Tuning either the cue or the feedback strength might have altered EVC behavior, however, we kept them identical to the settings in the other models for better comparison.

We applied the cue to the frontal modules on the basis of theoretical considerations of the nature of retro-cues. Unlike exogenous cues that highlight the location of a to-be-attended item in a bottom-up manner, endogenous cues as used in [26] require additional processing to trigger selection of the appropriate item from VWM. Therefore, we decided to apply the cue to the frontal modules, which are a proxy for high-level cortical areas capable of the required higher-level processing. With this setup, some feedback might be necessary to allow the propagation of this high-level information back to EVC. However, it is possible that top-down attentional gain acts on visual cortex directly as a result of processing the retro-cue. Therefore, we also tested whether applying the retro-cue to EVC directly would work to boost the cued representation. Specifically, we kept the model identical to the STSP model presented in the main text, but the cue was applied to EVC instead of frontal modules. As a result, the cued representation was deleted from EVC, as is evident by the lack of reactivation of the cued representation through the second perturbation (Figure B1c-d). We tested whether weaker cue strengths would work in giving the cued item an edge, but found that even the weakest cues of strength 0.01 already slightly degraded the decoding accuracy of the cued compared to the uncued item, and that the cued item got deleted for cue strengths equal to or larger than 1 (Figure B2).

**Figure B1:**
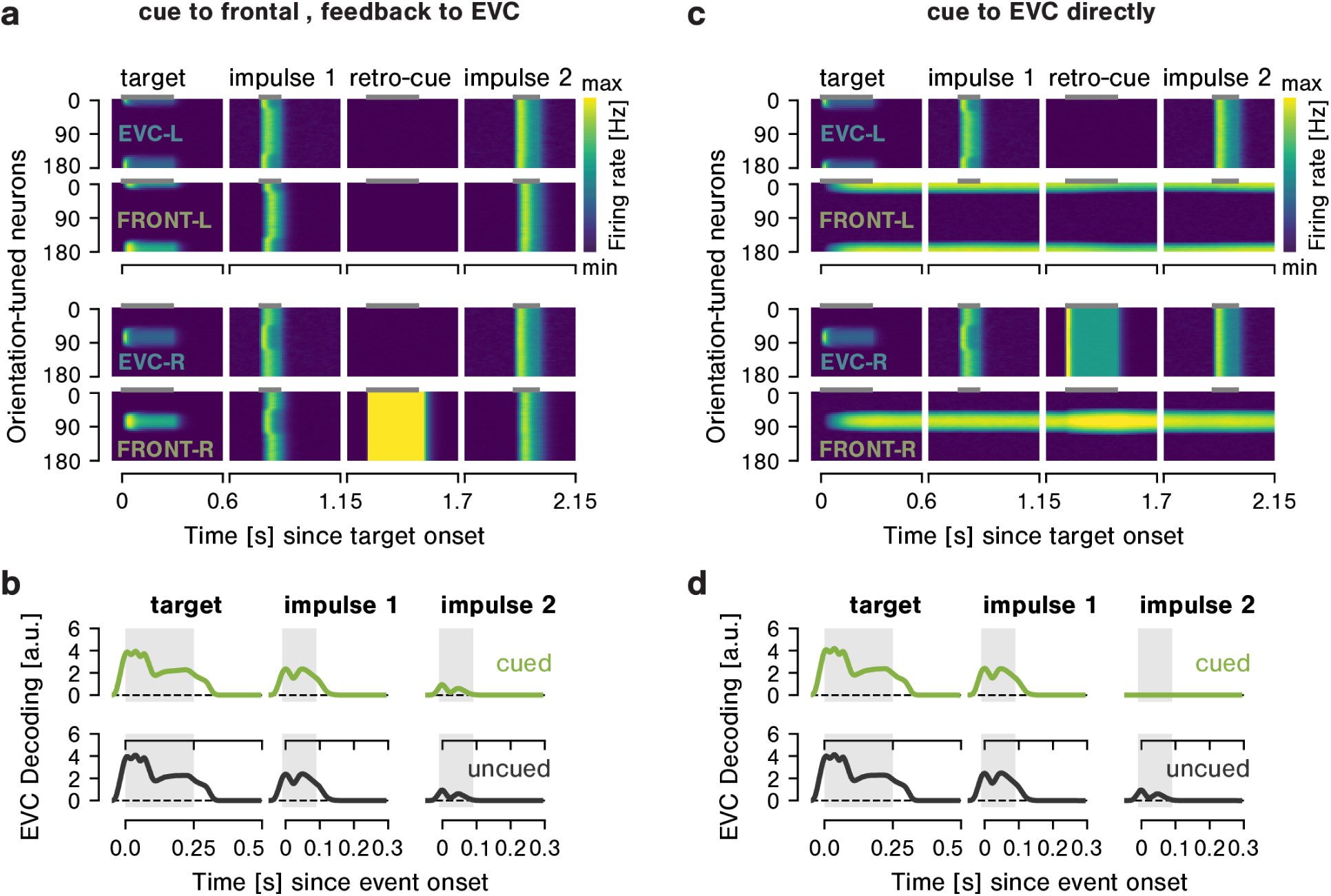
STSP model behaviour in the retro-cue task for different cueing strategies. **a** Neural responses in EVC and frontal module in response to stimulus, impulse, and cue presentation for an example trial. The cue is presented to the left frontal module, which is tuned to produce transient responses so as to avoid a memory forming anywhere but in EVC, and propagates back to EVC via an intact feedback connection. Due to the weak feedback connection, this does not elicit a population response in EVC. **b** Orientation decoding for the same model. Importantly, in response to the second impulse, there is no difference in decoding strength of the cued and the uncued items. **c-d** Same for an STSP model where the cue is applied to EVC directly. Importantly, as seen in **d**, the trace for the cued item is extinguished by the cue, as can be seen by the failure to decode the cued item during the second impulse.

**Figure B2:**
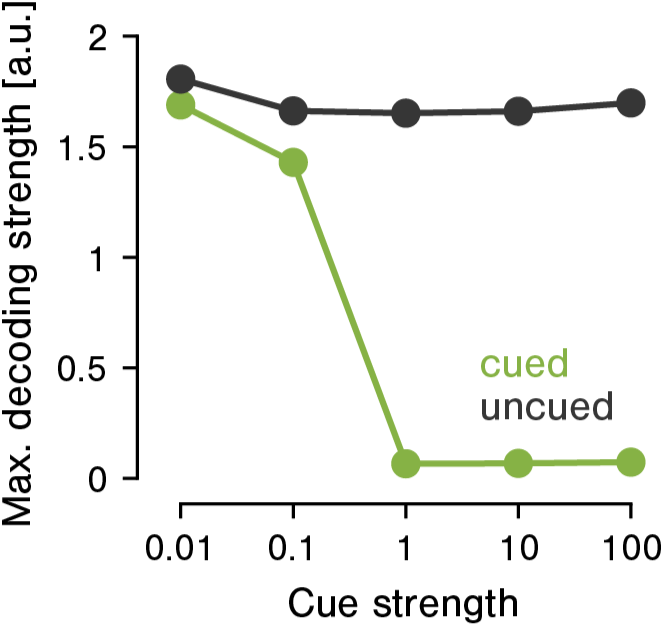
peak decoding strength of cued and uncued items during the second impulse as a function of cue strength. For cue strength => 1, Decoding of the cued item is extinguished.

## Appendix C

### Circuit Parameter Tuning

Neural circuits used in the frontal modules were tuned as in [78] to produce ring attractor dynamics. Circuits used in EVC were scaled down by a factor of 0.01 to produce transient dynamics. We used identical tuning for the STSP and feedback models. See Table C1 for an overview of tuning paramters. The last four parameters are specific to the EVC circuits in the STSP model.

**Table C1:** Parameter settings for EVC and frontal circuits.

| Parameter | EVC | Frontal |
| --- | --- | --- |
| $g_{EE}$ | 0.06 | 6 |
| $g_{EI}$ | 0.04 | 4 |
| $g_{IE}$ | 0.034 | 3.4 |
| $g_{II}$ | 0.0085 | 0.85 |
| $I_0^E$ | 0.002 | 0.2 |
| $I_0^I$ | 0.005 | 0.5 |
| $\sigma^E$ | 0.01 | 1 |
| $\sigma^I$ | 0.03 | 3 |
| $\tau^E$ | 20ms | 20ms |
| $\tau^I$ | 10ms | 10ms |
| $u_0$ | 0.2 | |
| $x_0$ | 1 | |
| $\tau_F$ | 1500ms | |
| $\tau_D$ | 200ms | |

## Appendix D

### Connectivity Matrices

Connectivity between modules within each model was mostly identical. As an exception, we scaled the feedforward connectivity between EVC and Frontal modules by a factor of 10 in the STSP model to account for the fact that firing rates were much lower in general due to the quickly depleting neurotransmitter resources. In addition, we omitted the feedback connection from frontal to EVC modules in the STSP model to isolate feedback vs. STSP as memory mechanisms in EVC. See Table D1 for an overview of connection strengths.

**Table D1:** Connection strengths among excitatory modules.

| Connection | structure | weight (STSP) | weight (FB) |
| --- | --- | --- | --- |
| **external inputs** |  |  |  |
| input (to EVC) | $N$-dimensional vector<br>(containing stimulus pattern) | 100 | 100 |
| impulse (to EVC) | $N$-dimensional vector<br>(vector of ones) | 0.5 | 0.5 |
| cue (to Frontal) | $N$-dimensional vector<br>(vector of ones) | 20 | 20 |
| **model connectivity** |  |  |  |
| $g_{ff}$ (EVC to Frontal,<br>feedforward) | $N * N$ identity matrix | 10 | 1 |
| $g_{EE}$ (Frontal to Frontal,<br>excitatory) | circulant matrix comprised of<br>$N$-dimensional von Mises<br>($loc = 0, \kappa = 5$) | 0.06 | 6 |
| $g_{mi}$ (Frontal to Frontal,<br>mutual inhibition) | $N * N$ matrix of ones | 1 | 1 |
| $g_{fb}$ (Frontal to EVC,<br>feedback) | $N * N$ identity matrix | 0 | 0.0001 |

